# Rapidly evolving homing CRISPR barcodes

**DOI:** 10.1101/055863

**Authors:** Reza Kalhor, Prashant Mali, George M. Church

## Abstract

We present here an approach for engineering evolving DNA barcodes in living cells. The methodology entails using a homing guide RNA (hgRNA) scaffold that directs the Cas9-hgRNA complex to target the DNA locus of the hgRNA itself. We show that this homing CRISPR-Cas9 system acts as an expressed genetic barcode that diversifies its sequence and that the rate of diversification can be controlled in cultured cells. We further evaluate these barcodes in cultured cell populations and show that they can record lineage history and and that their RNA can be assayed as single molecules *in situ*. This integrated approach will have wide ranging applications, such as in deep lineage tracing, cellular barcoding, molecular recording, dissecting cancer biology, and connectome mapping.

## Introduction

A single totipotent zygote has the remarkable ability to generate an entire multicellular organism. Methodologies to comprehensively map and modulate the parameters that govern this transformation will have far ranging impact on our understanding of human development and our ability to restore normal function in damaged or diseased tissues. One of these parameters that can provide important insights into developmental processes is the lineage history of cells (Sulston et al. 1983; Kretzschmar and Watt 2012). Contemporary lineage-tracing approaches, however, do not readily scale to model organisms, such as mice, that are most relevant to human development (Weisblat et al. 1978; Walsh and Cepko 1992; Dymecki and Tomasiewicz 1998; Lu et al. 2011; Porter et al. 2014). Precise mapping of lineage history in these organisms may be facilitated by combining modern genome engineering and DNA sequencing technologies (Mali, Esvelt, et al. 2013; Church et al. 2014; Lee et al. 2014): if every cell in an organism contained a unique and easily retrievable DNA sequence - a barcode - that encompassed its lineage relationship with other cells, this barcode could be probed to delineate the precise lineage history of all cells in the organism.

To this end, we propose here the concept of evolving genetic barcodes that are embedded in cells and change their genetic signature progressively over time (Figure 1). In a generalizable manifestation, this approach entails an array of genomically integrated sites that are stochastically targeted by a nuclease. During each cell cycle, the nuclease targets a random subset of these sites where the process of non-homologous end joining (NHEJ) introduces insertions, deletions, or other mutations, leading to a unique sequence that is related to its parent sequence and may further evolve in subsequent rounds. At the end of the developmental process, single-cell assaying technologies can be applied to each cell to decipher its unique barcode - the sequences of its nuclease site array - and delineate its lineage history (Figure 1).

**Figure 1.**
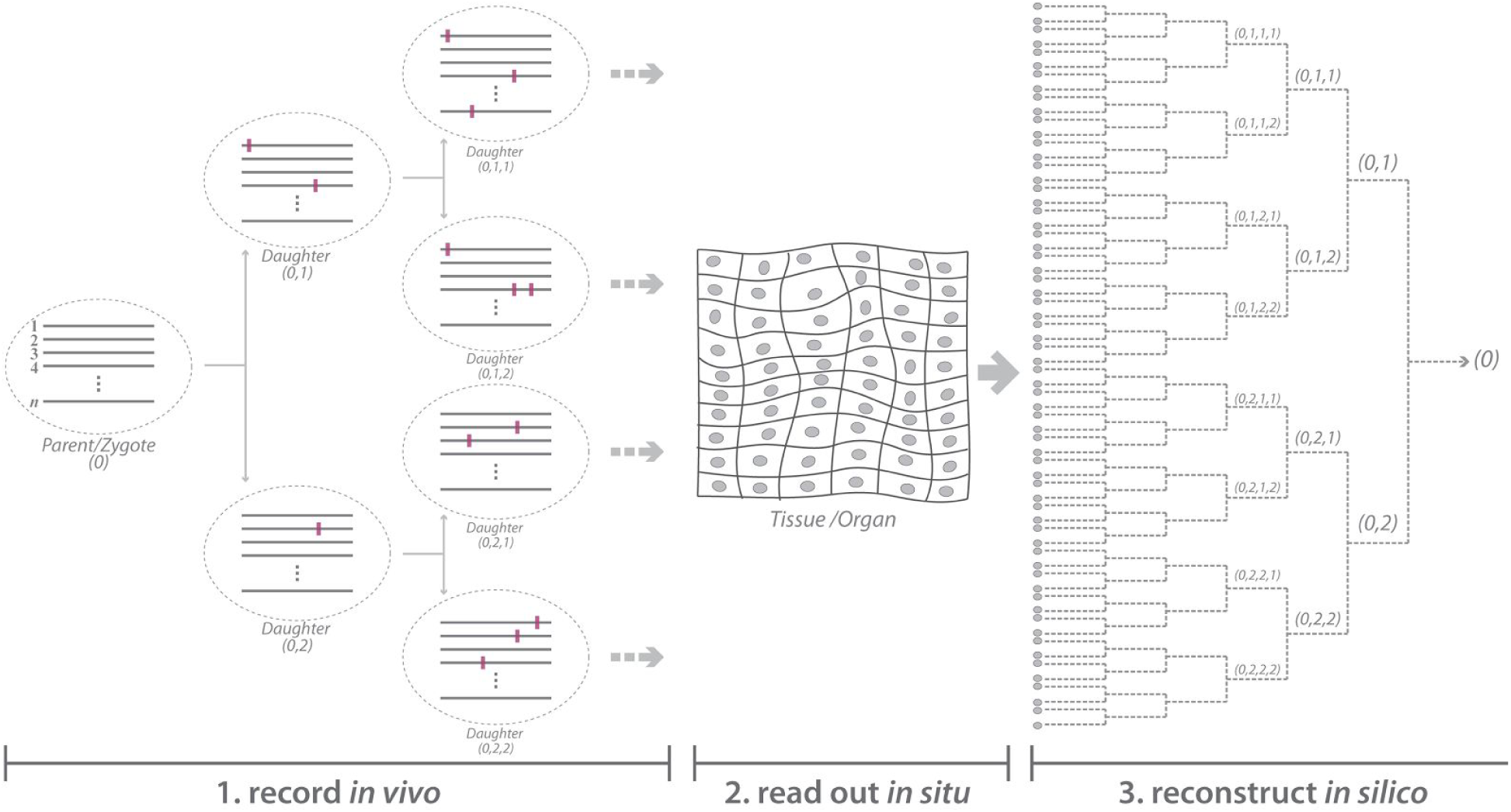
Schematic representation of a lineage tracing approach in multicellular eukaryotes using genome engineering and single cell DNA sequencing. (Left) Lineage recording during development. Array of *n* barcoding sites are represented by gray lines, cells are represented by dashed ovals, and mutations are represented by the purple rectangles. (Center) *In situ* read out of barcode sequences in tissues and organs. Gray ovals signify cell nuclei. (Right) *In silico* reconstruction of the lineage dendrogram based on the *in situ* sequencing readout of each cell.

Such an array of evolving barcodes can in theory fulfill the requirements of precise lineage tracing. However, for a practical implementation into an animal model such as mouse, it will have to meet several critical criteria. Specifically, it must create a total diversity commensurate with the total number of cells being targeted. If the diversity of possible mutations in each barcode element is ‘*m*’ and the number of independent array elements (i.e., without crosstalk) is ‘*n*’, then the system allows for creation of *m*^*n*^ possible signatures. Therefore, any system that generates higher values of *m* and *n* would be highly desirable. Furthermore, the system should continue to generate diversity throughout the development of the animal, and present tractable options for stable animal lines. Finally, it should be scalably readable at a single-cell level.

With above concepts in mind, we present homing guide RNAs (hgRNAs), a modified CRISPR-Cas9 system which targets the DNA locus of the guide RNA itself. We show that this simple system generates more diversity than canonical CRISPR-Cas9 (higher *m*). We further show that it is consistent with deployment in an array format with independently acting barcoding elements (higher *n*), and that the rate of diversification can be regulated to match the requisite pace for most model organisms. Additionally, we show that these barcodes are appropriate for lineage tracing applications *in vitro*, and their corresponding small RNAs can be assayed as single molecules *in situ*. We propose that these properties make hgRNAs a great candidate to integrate into animal models for barcoding and lineage tracing purposes.

## Results

The canonical CRISPR-Cas9 system can introduce mutations at a target locus via the process of non-homologous end joining (NHEJ). This system involves three components: a guide RNA (gRNA), the Cas9 protein, and a target site(Haurwitz et al. 2010; Horvath and Barrangou 2010; Jinek et al. 2012; Mali, Yang, et al. 2013a). The gRNA comprises a constant scaffold which establishes its interaction with the Cas9 protein and a variable spacer whose sequence determines the target site. To be digested by the Cas9:gRNA complex, in addition to matching the spacer sequence, the target site has to also include a specific protospacer adjacent motif (PAM) that is directly recognized by the Cas9 protein. As gRNAs do not have a PAM, loci that code for canonical gRNAs - or sgRNAs - are not targeted by the Cas9:gRNA complex despite containing the spacer sequence (Figure 2A). We sought to engineer a homing CRISPR system that directs Cas9:gRNA nuclease activity to the gRNA locus itself, thus simplifying the system by eliminating the target site and enabling retargeting and evolvability of barcodes (Figure 2B). To engineer a homing gRNA, we mutated the sequence immediately downstream of *S. pyogenes* gRNA spacer from ‘GUU’ to ‘GGG’ (Figure 2 C,D, Supplementary Figure 1A), so it matches the requisite ‘NGG’ PAM sequence of *S. pyogenes* Cas9. These bases are a part of a helix in the secondary structure of the canonical sgRNA (Figure 2C). To preserve the helix and minimize adverse structural impacts from our mutations, we also introduced compensatory mutations in the hybridizing nucleotides (Figure 2D). We then evaluated the functionality of this homing gRNA using an assay based on homologous recombination (Supplementary Figure 1B). In this assay, a ‘broken’ GFP gene is targeted by Cas9:gRNA complex in the presence of a repair template (Mali, Yang, et al. 2013b). Successful targeting of the broken GFP gene results in its repair through homologous recombination and the ensuing restoration of fluorescence can be detected. The results showed that our homing gRNA, or hgRNA, can digest a target sequence (Supplementary Figure 1B).

**Figure 2.**
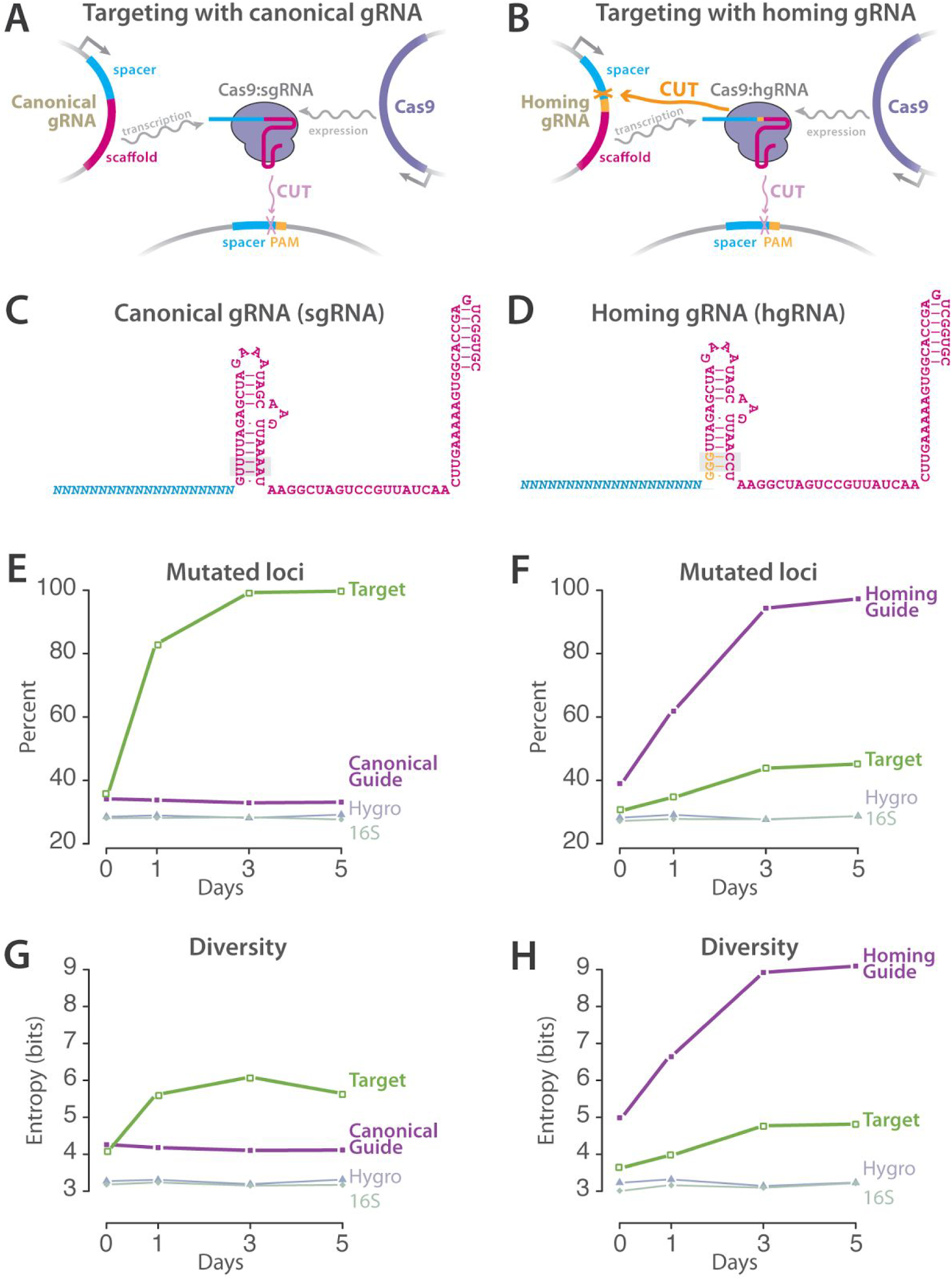
Comparison between standard and homing CRISPR-Cas9 systems. (A) Canonical CRISPR-Cas9 system, where Cas9 and sgRNA are expressed from their respective loci, form a complex and cut their target that matches both the spacer sequence of the gRNA and contains a PAM. (B) Homing CRISPR-Cas9 system, where Cas9-hgRNA complex, in addition to cutting their target sequence, also target the locus encoding the hgRNA itself. (C,D) Primary sequences and the predicted secondary structures of canonical and homing gRNAs. Positions that were mutated to derive the hgRNA are underlined. Orange bases mark the newly created PAM in hgRNA. Grey boxes mark all the positions where hgRNA is mutated compared to canonical sgRNA. (E,F) Accumulation of mutations in gRNA, target, a fragment of Hygromycin, and a fragment of the 16S ribosomal RNA loci upon Cas9 expression in cells with either canonical (panel E) or homing (panel F) versions of gRNA-A21. (G,H) Generated diversity in experiments of panels E and F measured as the Shannon entropy of the frequency vector of all variants that were observed in each condition. Data points are means (n = 2, biological replicates; s.e.m. small and not distinguishable on the plot scale).

To validate that homing gRNAs target their own locus as well, we created a HEK/293T clonal cell line genomically integrated with the humanized *S. pyogenes* Cas9 under an inducible Tet-On promoter (293T-iCas9 cells). We introduced both the hgRNA locus and its target into the genome of these cells using lentiviral integration, simulating the components of Figure 2B. As a control, we created the same system only with the canonical non-homing version of the gRNA, simulating the components of Figure 2A. We then induced Cas9 expression and harvested DNA samples from the cell population at various intervals after induction and sequenced the gRNA and target loci. The results showed cutting of the target locus by both the sgRNA and the hgRNA constructs (Figure 2E,F). sgRNA was more efficient in inducing mutations in the target locus; however only hgRNA induced mutations in its own DNA (Figure 2F, Supplementary Figure 1C). In fact, the hgRNA was as efficient in mutating its own DNA as the sgRNA was in mutating the target locus, both resulting in mutations in almost all cells five days after induction. These mutations were above background sequencing error rates that we measured by sequencing fragments of a similar size from the Hygromycin resistance gene upstream of the gRNA loci on the lentiviral backbone and the 16S rRNA gene on the genomic DNA of the cells. These results show that our homing gRNA, which is altered to contain a PAM, mutates the DNA locus encoding it.

We next compared the diversity generated by hgRNAs to that of sgRNAs. Two factors are important in considering this diversity: first, the number of different variants, and second, the frequency distribution of those variants. The ideal barcoding locus would generate a very high number of variants and all with an equal likelihood. As a proxy for both these factors, we measured the Shannon entropy of the frequencies of all the variants generated by both our hgRNA and its corresponding sgRNA in their gRNA and target loci (Figure 2 G,H). The results show that hgRNA can generate about 5 bits of diversity in its locus after Cas9 expression. This amount is a substantial improvement over the 2 bits generated by the sgRNA counterpart in its target and does not appear to come at a substantial cost to cell viability (Supplementary Figure 1D). The more diverse output, which is likely due to the evolution of hgRNAs past the first set of NHEJ products, suggest that hgRNAs are more suitable than sgRNAs for barcode generation.

The amount of diversity generated by a single hgRNA suggests that uniquely barcoding all neurons in a mouse brain requires an array of at least 6 hgRNAs per cell (see discussion). To assess whether other hgRNAs with a different spacer sequence show a similar behavior, we created six additional hgRNAs (B21, C21, D21, E21, F21, and G21) (Figure 3A, Supplementary Sequences). We assayed these hgRNAs in 293T-iCas9 cells (Figure 3B,C, Supplementary Table 1). The results showed that five of the six hgRNAs are highly active in targeting their parent loci and generate a similar amount of diversity, ranging from 5 to 6 bits. The hgRNA-B21 that showed a much lower activity level has a spacer with multiple ‘GG’ dinucleotides, a feature which has previously been shown to inhibit gRNA activity (Malina et al. 2015). These observations suggest that hgRNAs are generally functional irrespective of their spacer sequence and can thus operate as barcoding loci with minimal crosstalk between them.

**Figure 3.**
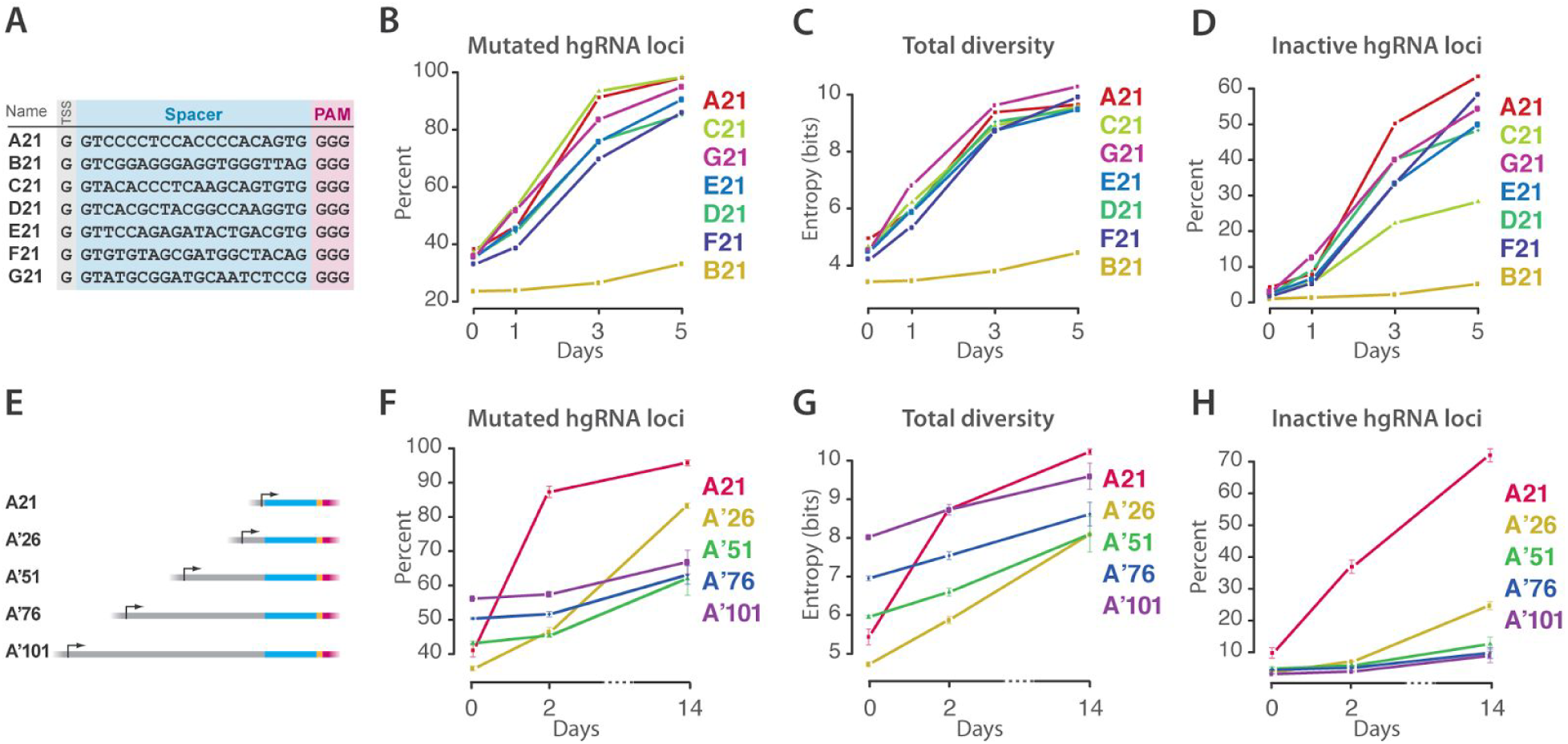
Performance of various hgRNAs. (A) The sequence of seven different hgRNAs. Transcription start site (TSS), spacer sequence, and PAM are marked with grey, blue, and pink boxes, respectively. (B,C,D) Cas9 is induced in cell lines with the hgRNAs from panel A integrated genomically. DNA samples harvested before (t=0 days) and at various points after induction are characterized by high-throughput sequencing to quantify mutations, functionality, and generated diversity for each hgRNA. Data points represent single replicates. See also Supplementary Table 1. (E) Design of four longer variants of hgRNA-A21. Stuffer sequences of 5, 30, 55, or 80 base pairs were added upstream of a spacer very similar to the A21 spacer to obtain the four increasingly lengthy A’ variants. (F,G,H) Cas9 is induced in cell lines with the hgRNAs from panel E integrated genomically. DNA samples harvested before (t = 0 days) and at various points after induction are characterized by high-throughput sequencing to quantify mutations and functionality of hgRNA loci. Data points are mean ± s.e.m. (n = 2, biological replicates).

Our first hgRNA set generated diversity for only a short time after induction of Cas9 expression before being inactivated (Figure 3D). While some were inactivated due to deletions that removed the PAM from the scaffold, others were rendered inactive by a truncation of their spacer below the 16-18 nucleotides necessary for Cas9-gRNA cleavage (Supplementary Table 1). As our hgRNA set had only 21 total bases between the RNA transcription start site and the scaffold, even small deletions in the spacer would led to truncated hgRNAs. We therefore sought to evaluate whether hgRNAs’ active lifespan can be prolonged by increasing their lengths. As such, based on hgRNA-A21, we created four variants that were 5, 30, 55, and 80 bases longer than hgRNA-A21 but had a similar initial spacer sequence (Figure 3E, Supplementary Figure 2, Supplementary Sequences). These hgRNAs were all active in our standard assay (Figure 3F, Supplementary Figure 2B), with the mutation and diversification rates decreasing with increasing hgRNA length (Figure 3F,G, Supplementary Figure 2B,C). Furthermore, unlike A21, these longer hgRNAs continued to generate diversity for a few weeks after induction (Figure 3H, Supplementary Figure 2D). These results show that hgRNA activity level can be regulated to accommodate developmental processes in the scale of weeks as well as those in the scale of days.

We further assessed whether hgRNAs can fulfill lineage tracing schemes similar to Figure 1. To simplify the experiment, instead of using multiple hgRNAs in a single parent cell and establishing the lineage relationship of its daughter cells, we used a single hgRNA in a parent cell population and attempted to decipher the lineage relationship of subpopulations that were derived from this parental population (Figure 4A,B). Specifically, we created one cell line with hgRNA-A21 (Figure 4A) and another with hgRNA-C21 (Figure 4B). From each of these parental lines, we established a first generation of daughter subpopulations usings ~100 cells and a brief Cas9 induction to generate diversity in the hgRNA locus. In a similar fashion, a second generation of daughter subpopulations was created from the first and a third from the second (Figure 4A,B). In the end, the hgRNA locus was sequenced in each subpopulation to profile its variants. The presence of non-shared variants between subpopulations was used as a measure of distance between them (Supplementary Table 2, Materials and Methods). Based on these distances, we clustered the second (Figure 4C, left and Figure 4D left) and third (Figure 4C, right and Figure 4D right) generations of subpopulations and found that their lineage relationship can be reconstituted from sequencing data. These results confirm that the diversity generated in hgRNA loci can be used to inform their lineage relationship.

**Figure 4.**
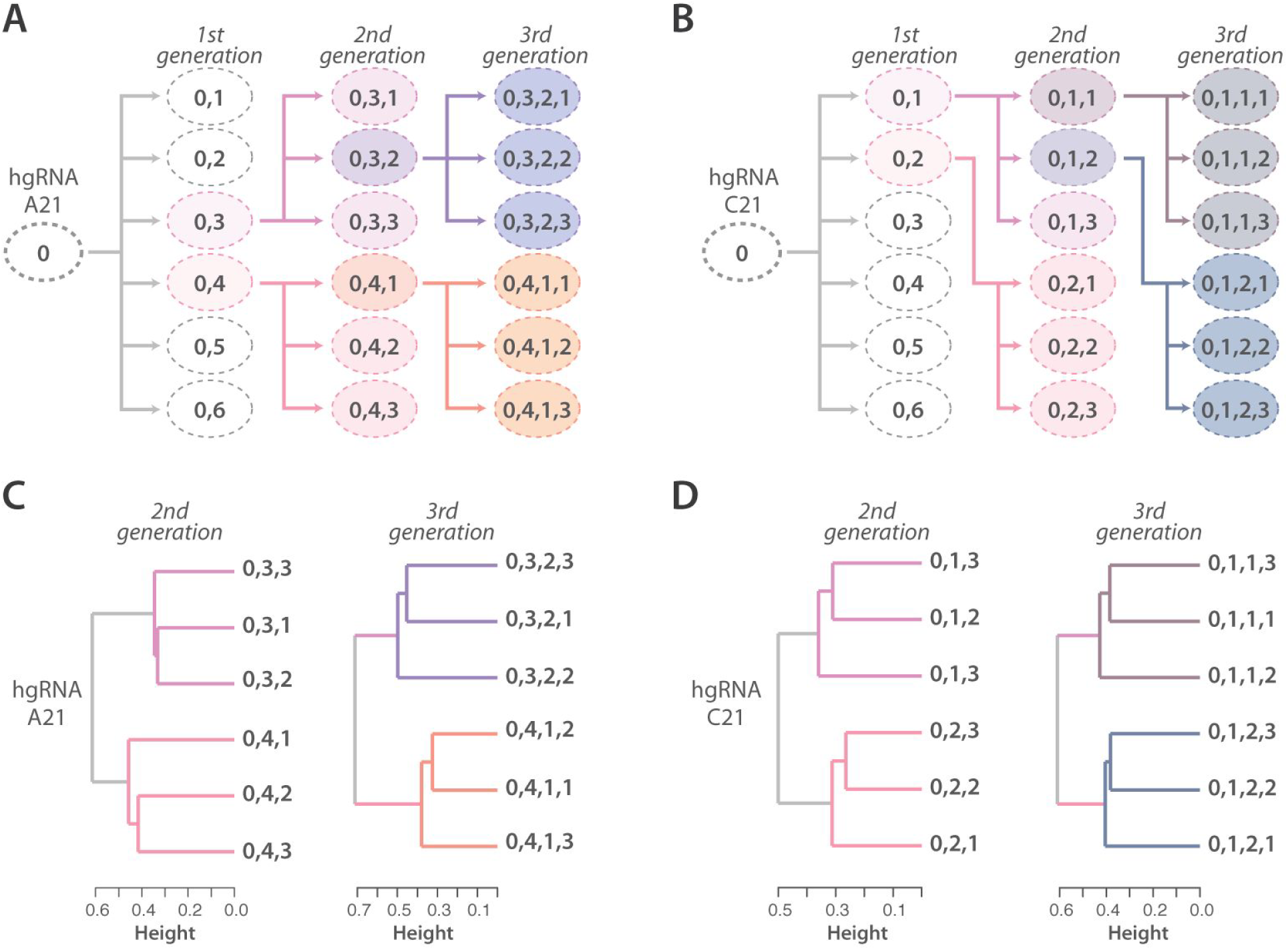
Lineage tracing in cultured cell populations. (A,B) Passaging scheme: cells carrying hgRNA-A21 (panel A) or hgRNA-C21 (panel B) were induced to express briefly Cas9, generating a parent population (‘0’) with a diverse array of hgRNA barcodes. As shown by arrows, the parent population (‘0’) was passaged into multiple first generation subpopulations (0,N), some of which were then further passaged into second and third generation subpopulations (0,N,N and 0,N,N,N respectively). The ovals represent populations and the numbers in the ovals represent the label of that line while indicating its relationship to the other lines. Each daughter subpopulation was seeded from about 100 cells that were briefly induced to express Cas9 immediately before seeding. (C,D) Clustering of subpopulations in panels A and B based on the hgRNA variants observed in each (Supplementary Table 2).

Finally, we turned our attention to barcode readout strategies, as retrievability is a requirement for effective lineage tracing. In this regard, *in situ* sequencing is a highly desirable method for barcode retrieval as it allows lineage information to be extracted without losing histological information such as position and cell type (Ke et al. 2013; Lee et al. 2014). One limitation of fluorescent *in situ* sequencing (FISSEQ) technologies is their short 20 to 30bp read length. Our approach is uniquely appropriate for retrieval with FISSEQ as the spacer sequence of the hgRNA loci where the barcode is generated is 20bp in length. However, available FISSEQ technologies probe arbitrary regions in a stochastic subfraction of longer transcripts in a cell; they face difficulty in both targeted sequencing and detection of smaller RNA molecules (Ke et al. 2013; Lee et al. 2014; Lee et al. 2015). We therefore assessed whether hgRNAs, which are small, can be probed in a targeted fashion using FISSEQ. FISSEQ can be divided into two stages: amplicon generation and amplicon sequencing. The difficulties associated with targeted probing of small RNA molecules relate to the amplicon generation step (Ke et al. 2013; Lee et al. 2014; Lee et al. 2015); thus, we used an assay to specifically address the amplification step for gRNAs (Figure 5A). We then executed the assay on representative hgRNA constructs with specific reverse-transcription (RT) primers that would target the barcoded region, and combined with the rolling-circle amplification (RCA) primer, allow for targeted amplification. We observed that RT and RCA primers that aim to amplify the entire hgRNA could not generate amplicons above the background level (Figure 5B, middle). We surmised that possible reasons for this failure were the strong secondary structure of the intervening hgRNA and the long distance (128bp) between the RT and RCA primers. We thus created a second version of the hgRNA construct by inserting the RT primer binding site further upstream, inside the hgRNA scaffold at positions previously shown tolerant of insertions (Mali, Aach, et al. 2013; Konermann et al. 2015). This new arrangement, resulted in robust *in situ* amplification and detection of the gRNA spacer region in a target-specific fashion (Figure 5B, right). These results address the challenges associated with *in situ* amplification step of gRNAs and hgRNAs and make it possible to read out hgRNA barcodes using *in situ* sequencing.

**Figure 5.**
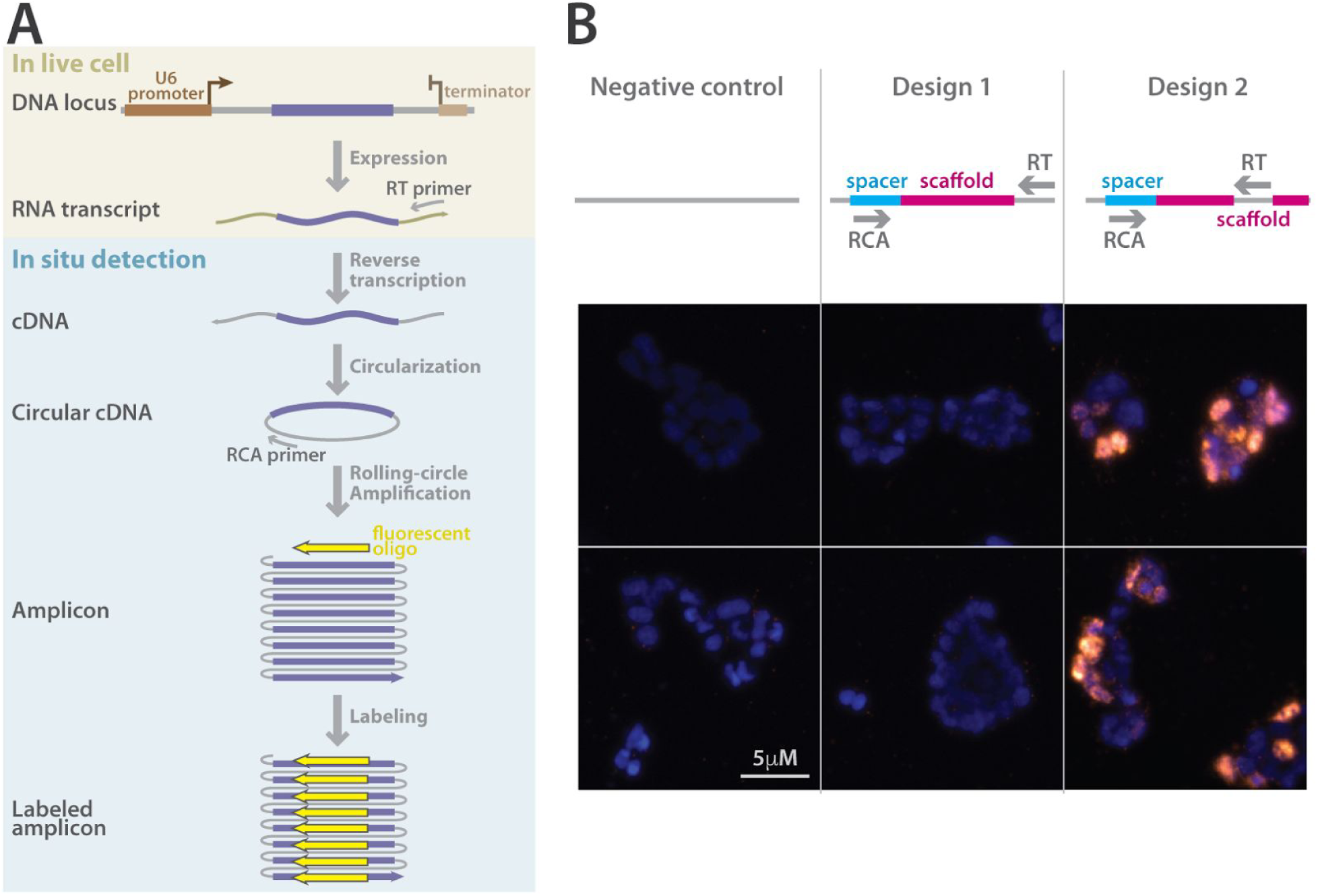
Target-specific *in situ* detection of gRNA molecules expressed by RNA polymerase III. (A) A schematic of our *in situ* amplification and detection assay based on FISSEQ. A DNA locus expressing a gRNA (purple) under the U6 promoter (brown) is introduced into cells. The construct also contains designed primer binding sites both downstream and upstream of the gRNA in grey-colored regions. A U6 terminator (light brown) is placed after the second primer binding region. Cells containing this locus, and thus expressing its RNA transcript, are fixed for in situ amplification and detection. In the fixed cells, the RNA transcript is reverse-transcribed using a locus-specific RT (reverse-transcription) primer to obtain a cDNA which is then circularized. The circular cDNA is amplified by the rolling circle amplification (RCA) using a second locus-specific RCA primer, producing a concatemerized amplicon that is confined to a small space in the hydrogel matrix of the experiment. The amplicon is then labeled by a fluorescent oligonucleotide. (B) Results of target-specific in situ amplification and detection for two different gRNA constructs and a negative control. The schematic on top shows the position of the reverse-transcription (RT) primer in each design. The bottom panels show a representative field of view from each experimental replicate for each construct. Amplicons are labeled with Cy5 (orange) and nuclei are labeled with DAPI (blue). The amplicon is only detectable in cell transfected with the Design 2 constructs, whereas Design 1 only shows very few labeled amplicons, at a level similar to the false positive amplicons in the negative control.

## Discussion

Recent reports also demonstrate the utility of nuclease mediated diversity generation in biological systems (Junker et al. 2016; Perli et al. 2016; McKenna et al. 2016 May 26). For instance, one study utilizes canonical gRNA-target pairs with decreasing affinities for lineage tracing in zebrafish (McKenna et al. 2016 May 26), and another uses homing CRISPR systems for recording analogue cellular signals (Perli et al. 2016). We apply the homing CRISPR system to engineer evolving barcodes for barcoding and lineage tracing purposes. We evaluate several important parameters of this system that reflect on its potential for barcoding and lineage tracing applications similar to Figure 1. Namely, we demonstrate that homing gRNAs generate more than eight times the diversity that canonical gRNAs generate (2^2^ versus 2^5^) (Figure 2G,H). We show that different hgRNA spacer sequences are functional (Figure 3C), providing a straightforward way to creating an array of these elements without crosstalk between them while each element is uniquely distinguishable. Additionally, we show that the diversification rate of hgRNAs can be adjusted (Figure 3G) to fit what may be necessary for specific applications or model organisms. We also show, in principle, application of multiple hgRNAs to tracing lineage in cell populations (Figure 4).

It is perhaps instructive to estimate the minimum number of barcoding elements that would yield a useful animal model. A homing gRNA locus is capable of storing about 5 bits of information which is enough to distinguish 2^5^ = 32 different states. Accordingly, uniquely barcoding the roughly 12 billion cells in a mouse will require at least 7 such hgRNA loci per cell ((2^5^)^7^ > 12 × 10^9^). Uniquely barcoding the estimated 75 million neurons in a mouse brain will require at least 6 such hgRNA loci per neuron ((2^5^)^6^ > 75 × 10^6^). While the actual number of barcoding loci needed in practice will depend on the tolerance of an experiment to duplicate barcodes, these estimates suggest that adequate barcoding of mouse neurons, which is essential for some of the proposed brain mapping projects in the brain initiative (Adam H. Marblestone et al. 2013; A. H. Marblestone et al. 2013; Church et al. 2014), may be within reach of our current strategy.

We further assess the feasibility of reading out small gRNA-based barcodes using current *in situ* sequencing technologies. We find that the gRNA backbone presents limitations for *in situ* amplification, a prerequisite of *in situ* sequencing, likely due to long amplicon length and RNA secondary structure. By inserting the reverse transcription primer site into the scaffold, we have created a modified version of hgRNA in which the barcode region can be amplified *in situ*. This scaffold modification enables targeted *in situ* sequencing of gRNAs and thus single-cell *in situ* probing of barcodes.

An important strength of this hgRNA-based barcoding system lies in its potential for establishing a stable transgenic line in a model organism, such as mouse. hgRNA barcoding elements are single-component genetic loci (Figure 2B). Canonical sgRNAs, in contrast, require paired gRNA and target elements (Figure 2A) to be integrated into model organisms, a problem that becomes increasingly complex with the requirement for arrays of multiple independent barcoding elements. Furthermore, while hgRNAs are tolerant of mutations in their spacer region, canonical sgRNAs are susceptible to inactivation by mutations in their gRNA spacer or the target protospacer, presenting further challenges for creating stable transgenic lines. Finally, compared to non-CRISPR nuclease systems, a barcoding model organism with hgRNAs offers all the flexibility and versatility of Cas9 without a need to recreate the transgenic line that carries the hgRNA array: any modified or tissue specific Cas9 can be delivered by viral vectors or through crossing with a transgenic animal.

We also note three key limitations in our current implementation: first, the limited duration of evolvability for the fastest evolving hgRNAs and the still limited diversity generated by each element; we hope that the use of improved and orthogonal inducible systems (Platt et al. 2014), coupled with large arrays of evolving barcodes, as well as the use of molecules that modulate NHEJ outcomes (such as end processing enzymes, polymerases and terminal transferases (Certo et al. 2011)) or base-editing Cas9s (Komor et al. 2016)can substantially enhance both the durability of hgRNAs and the amount of diversity they generate. Second, the current limitations associated with *in situ* read out in general. In this regard, enhanced imaging capabilities (Liu and Keller 2016) and automation of *in situ* sequencing can pave the way for ultra-dense readout of cellular barcodes. Third, the exact effect of integrating an array of nuclease sites into to the genome on cell viability and developmental pathways themselves is unclear. Substantial work is required to determine if and to what extent such effects exist and how they can be mitigated.

In conclusion,*in vivo* barcode array strategies combined with *in situ* readout, similar to the one outlined here, can leverage the remarkable diversity of DNA sequences to study biological systems and will have broad applications in developmental biology, molecular recording (Farzadfard and Lu 2014), cancer biology, and in mapping neural connectivity (Zador et al. 2012; A. H. Marblestone et al. 2013; Naik et al. 2014; Peikon et al. 2014).

## Materials and Methods

### Vector construction

The Cas9 vectors used in the study are based on earlier published work (Yang et al. 2013; Mali, Yang, et al. 2013b). The hgRNA vectors and the sgRNA counterpart of hgRNA-A21 were constructed by incorporating corresponding gBlock (IDT DNA) synthesized DNA fragments (spacers and scaffolds) into the pLKO.1 lentiviral backbone (MISSION shRNA vectors via SIGMA) which was modified to carry Hygromycin B resistance. The target locus of sgRNA-A21 and hgRNA-A21 were constructed by incorporating corresponding gBlock (IDT DNA) synthesized DNA fragment into the pLKO.1 lentiviral backbone (MISSION shRNA vectors via SIGMA) which was modified to carry Blasticidin resistance. This target locus was designed such that its sequence is identical to that of sgRNA-A21 in the region that is subjected to sequencing, however, it cannot act as a hgRNA because it lacks the U6 promoter and contains only a truncated scaffold. The sequence of these inserts is available in supplementary material.

### HEK/293T cells with inducible *S. pyogenes* Cas9

Humanized *S. pyogenes* Cas9 (hCas9) under the control of a Tet-On 3G inducible promoter and carrying the puromycin resistance gene was genomically integrated in Human Embryonic Kidney 293T cells (HEK/293T, ATCC CRL-11268) using a PiggyBac transposon system. Multiple clonal lines were derived from the transduced population and doxycycline-induced expression of hCas9 was measured in each line using reverse-transcription followed by quantitative PCR. The best line showed low baseline levels of hCas9 expression and about 300 fold enrichment of hCas9 upon induction (data not shown). This line was used in all ensuing experiments and will be referred to as 293T-iCas9. These cells were cultured on poly-D-lysine coated surfaces and in DMEM with 10% Fetal Bovine Serum and 1µg/ml Puromycin in all experiments.

### Lentivirus production

Lentiviruses were packaged in HEK/293T cells using a second generation system with VSV.G as the envelope protein. Viral particles were purified using polyethylene glycol precipitation and resuspension in PBS. They were stored at −80C until use.

### Transduction of 293T-iCas9 cells with lentiviral vectors carrying hgRNAs

293T-iCas9 cells were grown to 70% confluency at which point they were infected with 0.3-0.5 MOI of lentiviral particles in presence of 6µg/ml polybrene. About 3 days after infection, cells were placed under selection with 200µg/ml Hygromycin B. Cells were retained under selection for at least 1 week before any experiments to assure genomic integration. In all cases, loss of about half the entire cell population after selection was used to confirm single infection with lentiviruses. Cells were maintained under Hygromycin B selection throughout subsequent experiments.

### Double transduction of 293T-iCas9 cells with lentiviral vectors carrying standard or homing A21 guide RNA with and lentiviral vectors carrying their target

For the experiments where both the guide RNA and its target were expressed (Figure 2), 293T-iCas9 cells were grown to 70% confluency at which point they were infected either with sgRNA-A21-Hygromycin and A21-Target-Blasticidin (Figure 2 A,C,E,G) or with hgRNA-A21-Hygromycin and A21-Target-Blasticidin (Figure 2 B,D,F,H) with 0.3-0.5 MOI of lentiviral particles in presence of 6µg/ml polybrene. 3 days after infection, cells were placed under double selection with 200µg/ml Hygromycin B and 10µg/ml Blasticidin. Cells were retained under double selection for at least 1 week before any experiments to assure genomic integration. Cells were maintained under Hygromycin B and Blasticidin selection throughout subsequent experiments.

### Induction of hCas9 in cells with hgRNAs

For each cell line transduced with a hgRNA construct, a sample was harvested before induction to represent the state of the non-induced population. Cells were then induced with 2µg/ml doxycycline. At various time points after induction, as indicated for each experiment, a sample of the cells was harvested during a passage to represent different time points after induction. In one experiment, multiple samples were harvested from of a non-induced cell line at various time points (Supplementary Figure 1C).

### Lineage tracing in cultured cell populations

A293T-iCas9 cell line, carrying either hgRNA-A21 or hgRNA-C21, was subjected to two hours of induction with doxycycline to generated limited initial diversity in the hgRNA loci, thus creating each founder cell population, here referred to as ‘0’. After doxycycline was removed, about 100 cells from each ‘0’ populations were used to seed six subpopulations, ‘0,1’ through ‘0,6’, in 6.5mm wells. In the course of about ten days, each subpopulation was expanded into larger wells and a sample was taken for sequencing analysis. Two of the subpopulations were then randomly selected for further passaging. About 100 cells from each selected subpopulations were induced with doxycycline for two hours and used to seed the next group of daughter subpopulations, e.g., ‘0,4,1’ through ‘0,4,3’. This passaging scheme was repeated for one more round to create the third generation of daughter subpopulations (Figure 4a,b).

### High-throughput DNA sequencing

Genomic DNA was extracted from each sample using the Qiagen DNAeasy Blood&Tissue kit. To amplify gRNA loci (canonical or homing) the following primer pair was used:

SBS3_Guide_F acactctttccctacacgacgctcttccgatct atggactatcatatgcttaccgt

SBS9_Guide_R tgactggagttcagacgtgtgctcttccgatct ttcaagttgataacggactagc

These primers amplify a fragment starting 81bp upstream of the transcription start site for the U6 promoter and ending 55bp after the start of gRNA scaffold. The total fragment length varies for various hgRNA constructs, but in its shortest form (e.g., A21) is 157bp.

To amplify the target locus the following primer pair was used:

SBS3_Target_F acactctttccctacacgacgctcttccgatct aagaggatggtgcagcaaccaag

SBS9_Target_R tgactggagttcagacgtgtgctcttccgatct tcaatctgacaggtgcctctcac

These primers amplify a fragment starting 82bp upstream of the spacer sequence and ending 53bp after the PAM site. The total fragment length is 158bp.

To amplify the Hygromycin B resistance locus as a control, the following primer pair was used:

SBS3_Hyg_F acactctttccctacacgacgctcttccgatct gtcgatgcgacgcaatcgtc

SBS9_Hyg_R tgactggagttcagacgtgtgctcttccgatct ttcctttgccctcggacgag

These primers amplify a fragment starting 881bp downstream of the Hygromycin gene start codon and ending just before the stop codon - 1032bp after the start codon. The total fragment length is 152bp.

To amplify a part of the 16S rRNA locus as another control the following primer pair was used:

SBS3_16S_F acactctttccctacacgacgctcttccgatct atgcatgtctgagtacgcac

SBS9_16S_R tgactggagttcagacgtgtgctcttccgatct ccgaggttatctagagtcac

These primers amplify a 259bp fragment from the human 16S rRNA locus.

PCR reactions with above primer pairs were carried out in a real-time setting and stopped in mid-exponential phase, typically about 20 cycles. To add the complete Illumina sequencing adaptors, this first PCR product was diluted and used as a template for a second PCR reaction with NEBNext Dual Indexing Primers, with each sample receiving a different index. Once again, PCR was carried out in a real-time setting and stopped in mid-exponential phase, which was after about 15 cycles in all cases. Samples were then combined and sequenced using Illumina MiSeq with reagent kit v3. Sequencing was done in one direction, starting from the forward (F) primer in each sample and for 170bp on average, but longer for libraries with longer hgRNA constructs.

### High-throughput DNA sequencing analysis

For each gRNA locus that was subjected to sequencing, only the fragment starting 4bp before the expected transcription start site for U6 promoter and ending 32bp into the scaffold was considered during all below analyses. For hgRNA-A21, for instance, this fragment is 57bp in total length. For the A21 Target locus only the 57bp fragment exactly equivalent to that of the hgRNA-A21 or sgRNA-A21 was considered. For the Hygromycin and 16S rRNA loci, which acted as controls, only a 57bp fragment with the same relative start and end positions in the sequencing read as hgRNA-A21 were considered to control for potential positional variations in sequencing accuracy.

The frequency of each hgRNA variant or mutant, which contained at least one mismatch, deletion, or insertion compared to the sequence of the original hgRNA, was determined for each sample. For the 57bp fragment considered for hgRNA-A21, sgRNA-A21, Hygromycin and 16S rRNA loci (Figure 2 E,F), an average sequencing error rate of 0.0064, characterized under standard conditions (Schirmer et al. 2015), results in 30% of all fragments to appear as mutants [1 - (1-0.0064)^57^]. This rate is in agreement with our observed values for Hygromycin and 16S rRNA loci and shows that the observed background mutant levels are largely a result of sequencing error. For longer construct, hgRNAs A’26, A’51, A’76, A’101, fragments of 62, 87, 112, and 137bp were considered in the analysis (Figure 3F,G,H). Considering an average sequencing error rate of 0.0064, the expected background mutant levels for these longer fragments would be 33%, 43%, 51%, 59% respectively. These background levels are also in agreement with baseline mutant fractions observed before Cas9 induction in Figure 3F.

The diversity of the hgRNA library that was produced as the result of Cas9 expression is represented by the Shannon Entropy of the vector of all variants’ frequencies in each condition. Because sequencing error produces some “apparent” diversity in each library, as a measure of true diversity we used the difference between the observed diversities after Cas9 induction and the diversity observed before induction in the same experiment. Separate experiments confirmed that mutation levels in non-induced samples remain steady in the course of our experimental times (Supplementary Figure 1C).

To obtain fast alignment of a large number of reads to a short template we used two-step approach. First, we aligned all reads to their expected template using blat, which was run on an in-house server. Blat helped determine insertions and deletions, while keeping mismatches intact. In cases where blat results indicated a deletion that was partially or entirely overlapping with an insertion, we used a dynamic programming algorithm with a match score of 5 and mismatch and deletion scores of 0 to optimize the alignment further. Based on the alignment results, we annotated the spacer, the PAM, and the sequenced portions of the promoter and scaffold from each sequenced hgRNA. A sequenced hgRNA was deemed functional, or capable of re-cutting itself, if it had a PAM and a functional spacer, as well as correct promoter and scaffold. A promoter was annotated as correct if 80% of its sequenced and non-primer overlapping bases correctly matched the consensus promoter. The scaffold was annotated as correct if 90% of its sequenced and non-primer overlapping bases, excluding the PAM bases, correctly matched the expected scaffold. PAM was considered as correct if it matched the NGG sequence, NHG, or NGH (it was noticed that non-NGG PAMs, such as NHG and NGH, showed substantial activity). The spacer was considered functional when it was longer than 17 bases (deletions often lead to inactive hgRNAs with shortened spacers if the distance between the U6 transcription start site and PAM is short). For lineage tracing in cultured cell, first all hgRNA variants that were present in each subpopulation were determined. For that, the observed frequency of each gRNA variant was measured. Any variant with an observed frequency of at least 0.01% in a subpopulation was considered present in that subpopulation (setting this cutoff to as high as 1% and as low as 0.001% did not change downstream results), otherwise it was considered absent. As such, for each subpopulation, a binary vector was obtained with 1 for present hgRNA variants and 0 for absent ones. A binary distance matrix was constructed from these vectors (Supplementary Table 2). The vectors for the subpopulations at the same level were clustered hierarchically with a complete agglomeration strategy (Becker et al. 1988).

### In situ amplification and detection

HEK/293T cells were seeded at 10,000 per well in 96-well polystyrene dishes coated with poly-D-lysine. 12 hours later, each well was transfected with 100ng of plasmid a plasmid DNA packaged with 0.5µL of Lipofectamin 2000 reagent (ThermoFisher Scientific) accordingly to the manufacturer protocol. Positive samples received plasmids encoding for Design 1 or Design 2 constructs. Negative control samples received a GFP plasmid. 24 hours after transfection, cell were subjected to in situ amplification and detection of the gRNA transcripts.

In situ detection was carried out according to the previously described sequencing in situ sequencing protocol (Lee et al. 2014; Lee et al. 2015). In brief, cells were fixed using formalin and permeabilized. Reverse-transcription was then carried out using a target-specific primer (5P-tcttctgaaccagactcttgtcattggaaagttggtataagacaacagtg) in present of aminoallyl-dUTP. Nascent cDNA strands were crosslinked by treatment with BS(PEG)9 (ThermoFisher Scientific) and RNA was degraded by RNaseA and RNaseH treatment. cDNA was circularized using CircLigaseII (Epicentre). Rolling circle amplification (RCA) was carried out with Phi29 polymerase using a target-specific primer (ggtggagcaattccacaacac) overnight in presence of aminoallyl-dUTP. Nascent amplicons or ‘rolonies’ were crosslinked by treatment with BS(PEG)9. Target amplicons were labeled with a fluorescent target-specific detection probe (5Cy5-tcttctgaaccagactcttgt) which recognizes the reverse-transcription primer and nuclei were stained with DAPI. Samples were imaged with a Zeiss Observer.Z1 inverted microscope using a 20X magnification objective in the DAPI and Cy5 channels.

## Accession Codes

The sequencing files are available in the Sequence Read Archive with the identifier SRP092492.

## Acknowledgements

The authors would like to acknowledge Wei Leong Chew, John Aach, Susan Byrne, Evan Daugharthy, Thomas Ferrante, Je Hyuk Lee, Mark Moosburner, Ian Peikon, Henry Lee, Alex Ng, Javier Fernandez Juarez, Adam Marblestone, Alex Chavez, Yoav Mayshar, Jonathan Scheiman, Kian Kalhor, Ting Wu, Jay Shendure, and Tim Lu for helpful comments or discussions and the Biopolymers Facility at HMS for technical assistance. This work has been supported by funding from NIH grants MH103910, HG005550 and the Intelligence Advanced Research Projects Activity (IARPA) via Department of Interior/Interior Business Center (DoI/IBC) contract number D16PC00008, and UCSD new faculty startup funds (PM).

## References

Becker RA, Chambers JM, Wilks AR. 1988. The new S language: a programming environment for data analysis and graphics. Chapman & Hall/CRC.

Certo MT, Ryu BY, Annis JE, Garibov M, Jarjour J, Rawlings DJ, Scharenberg AM. 2011. Tracking genome engineering outcome at individual DNA breakpoints. Nat. Methods 8:671–676.

Church GM, Marblestone AH, Kalhor R. 2014. Rosetta Brain. In: Marcus G, Freeman J, editors. The Future of the Brain: Essays by the World’s Leading Neuroscientists. Princeton: Princeton University Press. p. 50–66.

Dymecki SM, Tomasiewicz H. 1998. Using Flp-recombinase to characterize expansion of Wnt1-expressing neural progenitors in the mouse. Dev. Biol. 201:57–65.

Farzadfard F, Lu TK. 2014. Synthetic biology. Genomically encoded analog memory with precise in vivo DNA writing in living cell populations. Science 346:1256272.

Haurwitz RE, Jinek M, Wiedenheft B, Zhou K, Doudna JA. 2010. Sequence- and structure-specific RNA processing by a CRISPR endonuclease. Science 329:1355–1358.

Horvath P, Barrangou R. 2010. CRISPR/Cas, the Immune System of Bacteria and Archaea. Science 327:167–170.

Jinek M, Chylinski K, Fonfara I, Hauer M, Doudna JA, Charpentier E. 2012. A programmable dual-RNA-guided DNA endonuclease in adaptive bacterial immunity. Science 337:816–821.

Junker JP, Spanjaard B, Peterson-Maduro J, Alemany A, Hu B, Florescu M, van Oudenaarden A. 2016. Massively parallel whole-organism lineage tracing using CRISPR/Cas9 induced genetic scars. bioRxiv:056499. [accessed 2016 Sep 19]. http://biorxiv.org/content/early/2016/06/01/056499

Ke R, Mignardi M, Pacureanu A, Svedlund J, Botling J, Wählby C, Nilsson M. 2013. In situ sequencing for RNA analysis in preserved tissue and cells. Nat. Methods 10:857–860.

Komor AC, Kim YB, Packer MS, Zuris JA, Liu DR. 2016. Programmable editing of a target base in genomic DNA without double-stranded DNA cleavage. Nature 533:420–424.

Konermann S, Brigham MD, Trevino AE, Joung J, Abudayyeh OO, Barcena C, Hsu PD, Habib N, Gootenberg JS, Nishimasu H, et al. 2015. Genome-scale transcriptional activation by an engineered CRISPR-Cas9 complex. Nature 517:583–588.

Kretzschmar K, Watt FM. 2012. Lineage tracing. Cell 148:33–45.

Lee JH, Daugharthy ER, Scheiman J, Kalhor R, Ferrante TC, Terry R, Turczyk BM, Yang JL, Lee HS, Aach J, et al. 2015. Fluorescent in situ sequencing (FISSEQ) of RNA for gene expression profiling in intact cells and tissues. Nat. Protoc. 10:442–458.

Lee JH, Daugharthy ER, Scheiman J, Kalhor R, Yang JL, Ferrante TC, Terry R, Jeanty SSF, Li C, Amamoto R, et al. 2014. Highly multiplexed subcellular RNA sequencing in situ. Science 343:1360–1363.

Liu Z, Keller PJ. 2016. Emerging Imaging and Genomic Tools for Developmental Systems Biology. Dev. Cell 36:597–610.

Lu R, Neff NF, Quake SR, Weissman IL. 2011. Tracking single hematopoietic stem cells in vivo using high-throughput sequencing in conjunction with viral genetic barcoding. Nat. Biotechnol. 29:928–933.

Malina A, Cameron CJF, Robert F, Blanchette M, Dostie J, Pelletier J. 2015. PAM multiplicity marks genomic target sites as inhibitory to CRISPR-Cas9 editing. Nat. Commun. 6:10124.

Mali P, Aach J, Benjamin Stranges P, Esvelt KM, Moosburner M, Kosuri S, Yang L, Church GM. 2013. CAS9 transcriptional activators for target specificity screening and paired nickases for cooperative genome engineering. Nat. Biotechnol. 31:833–838. [accessed 2016 May 26]

Mali P, Esvelt KM, Church GM. 2013. Cas9 as a versatile tool for engineering biology. Nat. Methods 10:957–963.

Mali P, Yang L, Esvelt KM, Aach J, Guell M, DiCarlo JE, Norville JE, Church GM. 2013a. RNA-Guided Human Genome Engineering via Cas9. Science 339:823–826. [accessed 2016 May 26]

Mali P, Yang L, Esvelt KM, Aach J, Guell M, DiCarlo JE, Norville JE, Church GM. 2013b. RNA-Guided Human Genome Engineering via Cas9. Science 339:823–826. [accessed 2016 May 26]

Marblestone AH, Daugharthy ER, Kalhor R, Peikon ID, Kebschull JM, Shipman SL, Mishchenko Y, Lee JH, Dalrymple DA, Zamft BM, et al. 2013. Conneconomics: The Economics of Dense, Large-Scale, High-Resolution Neural Connectomics.

Marblestone AH, Zamft BM, Maguire YG, Shapiro MG, Cybulski TR, Glaser JI, Amodei D, Stranges PB, Kalhor R, Dalrymple DA, et al. 2013. Physical principles for scalable neural recording. Front. Comput. Neurosci. 7:137.

McKenna A, Findlay GM, Gagnon JA, Horwitz MS, Schier AF, Shendure J. 2016 May 26. Whole organism lineage tracing by combinatorial and cumulative genome editing. Science:aaf7907. [accessed 2016 May 26]

Naik SH, Schumacher TN, Perié L. 2014. Cellular barcoding: a technical appraisal. Exp. Hematol. 42:598–608.

Peikon ID, Gizatullina DI, Zador AM. 2014. In vivo generation of DNA sequence diversity for cellular barcoding. Nucleic Acids Res. 42:e127.

Perli S, Cui C, Lu TK. 2016. Continuous Genetic Recording with Self-Targeting CRISPR-Cas in Human Cells. bioRxiv:053058. [accessed 2016 May 27]. http://biorxiv.org/content/early/2016/05/20/053058

Platt RJ, Chen S, Zhou Y, Yim MJ, Swiech L, Kempton HR, Dahlman JE, Parnas O, Eisenhaure TM, Jovanovic M, et al. 2014. CRISPR-Cas9 knockin mice for genome editing and cancer modeling. Cell 159:440–455.

Porter SN, Baker LC, Mittelman D, Porteus MH. 2014. Lentiviral and targeted cellular barcoding reveals ongoing clonal dynamics of cell lines in vitro and in vivo. Genome Biol. 15:R75.

Schirmer M, Ijaz UZ, D’Amore R, Hall N, Sloan WT, Quince C. 2015. Insight into biases and sequencing errors for amplicon sequencing with the Illumina MiSeq platform. Nucleic Acids Res. 43:e37.

Sulston JE, Schierenberg E, White JG, Thomson JN. 1983. The embryonic cell lineage of the nematode Caenorhabditis elegans. Dev. Biol. 100:64–119.

Walsh C, Cepko CL. 1992. Widespread dispersion of neuronal clones across functional regions of the cerebral cortex. Science 255:434–440.

Weisblat DA, Sawyer RT, Stent GS. 1978. Cell lineage analysis by intracellular injection of a tracer enzyme. Science 202:1295–1298.

Yang L, Guell M, Byrne S, Yang JL, De Los Angeles A, Mali P, Aach J, Kim-Kiselak C, Briggs AW, Rios X, et al. 2013. Optimization of scarless human stem cell genome editing. Nucleic Acids Res. 41:9049–9061. [accessed 2016 May 26]

Zador AM, Dubnau J, Oyibo HK, Zhan H, Cao G, Peikon ID. 2012. Sequencing the connectome. PLoS Biol. 10:e1001411.

